# Correlations between metabolites in the synovial fluid and serum: a mouse injury study

**DOI:** 10.1101/2021.08.30.458234

**Authors:** Cameron W. Wallace, Brady Hislop, Alyssa K. Hahn, Ayten E. Erdogan, Priyanka P. Brahmachary, Ronald K. June

## Abstract

Osteoarthritis, the most common degenerative joint disease, occurs more frequently in joints that have sustained injury. Currently, osteoarthritis is diagnosed with imaging that finds radiographic changes after the disease has already progressed to multiple tissues. The primary objective of this study was to compare potential metabolomic biomarkers of joint injury between the synovial fluid and serum in a mouse model of post-traumatic osteoarthritis. The secondary objective was to gain insight into the pathophysiology of osteoarthritis by examining metabolomic profiles after joint injury. 12-week-old adult female C57BL/6 mice (n=12) were randomly assigned to control, day 1 post injury, or day 8 post injury groups. Randomly selected stifle (*i.e.,* knee) joints were placed into a non-invasive injury apparatus and subjected to a single dynamic axial compression causing anterior translation of the tibia relative to the femur to tear the anterior cruciate ligament. At days 1 and 8 post injury, serum was extracted then mice were immediately euthanized prior to synovial fluid collection. Metabolites were extracted and analyzed by liquid chromatography coupled to mass spectrometry. We detected ~2500 metabolites across serum and synovial fluid. Of these metabolites 179 were positively correlated and 51 were negatively correlated between synovial fluid and serum, indicating potential for the development of metabolomic biomarkers. Synovial fluid appeared to capture differences in metabolomic profiles between injured mice at both day 1 and 8 after injury whereas serum did not. However, synovial fluid and serum were distinct at both days 1 and 8 after injury. In the synovial fluid, pathways of interest across different time points mapped to amino acid synthesis and degradation, bupropion degradation, and the tRNA charging pathway. In the serum, notable pathways across time points were amino acid synthesis and degradation, the phospholipase pathway, and nicotine degradation. These results provide a rich picture of the injury response at early time points following traumatic joint injury. Furthermore, the correlations between synovial fluid and serum metabolites suggest that there is potential to gain insight into intra-articular pathophysiology through analysis of serum metabolites.

## Introduction

Osteoarthritis (OA) is one of the most common diseases in the United States, affecting more than 50 million people with a total annual cost exceeding $128 billion[1]. OA is a joint disease characterized by loss of cartilage, osteophyte formation, subchondral sclerosis, cysts, and joint space narrowing that manifest as pain and loss of function[2]. OA has long been considered a disease caused by degeneration of articular cartilage and bone. However, since the late 1990’s low-grade inflammation and other systemic factors have been increasingly recognized as relevant to the initiation and progression of OA[3, 4]. Joint injury results in post-traumatic osteoarthritis (PTOA). 50% of people with a diagnosed anterior cruciate ligament (ACL) tear or meniscus tear develop PTOA within 10-20 years[5]. Furthermore, imaging techniques can lack sensitivity and do not correlate well with symptoms[6]. PTOA is commonly diagnosed after significant tissue degeneration leading to current treatments being focused on pain management rather than prevention. These treatments include anti-inflammatories, cortisone injections, arthroscopic surgery, total joint arthroplasty, and sometimes surgical joint fusions. However, an emerging paradigm includes metabolic and structural changes within the joint that develop shortly after the initial injury thus providing a potential window for intervention[7].

Preclinical mouse models are important tools for studying PTOA. Common ways to induce PTOA in mice include ACL transection (ACLT), destabilization of the medial meniscus (DMM), and intra-articular injections of agents that induce joint degradation[8]. ACLT and DMM in mice can be challenging procedures and require incisions that prevent interpretation of early time points (e.g., < 10 days). Intra-articular injections, while less invasive, often induce only a single inflammatory pathway which while having major scientific value does not replicate *in vivo* pathophysiology. Non-invasive mouse models of PTOA have been developed that provide ACL rupture using tibial compression overload [8, 9]. This method offers the opportunity to evaluate early time points after injury and observe natural joint metabolomics induced by injury while simulating clinical ACL tears.

Because OA is a heterogenous disease, both diagnosis and treatment are complex[10]. The current American College of Rheumatology guidelines for diagnosing OA indicate that it is a clinical diagnosis confirmed with radiographs[11]. However, at the point of radiographic detection, irreversible changes have often already begun[12]. Additionally, there are currently no FDA-approved drugs to stop or reverse the progression of OA which motivates studying the early window immediately after joint injury.

Metabolites are small molecules that are generated from cellular processes. Metabolomic profiling is a technique that characterizes large numbers of metabolites (i.e., thousands) from biological samples including blood, urine, synovial fluid, and others via mass spectrometry and other advanced methods[13, 14]. Metabolomic profiles represent metabolic processes driven by cellular biochemistry and offer insight into phenotypical processes[15]. Thus, metabolomics has strong potential to detect metabolic changes early in the course of PTOA before symptomatic changes are detected which may be useful for clinical treatment including early diagnosis and phenotyping[15–17].

PTOA induces injury responses both within the joint and systemically[18]. The synovial fluid is in contact with most of the tissues of the joint, and metabolomic profiling of synovial fluid has strong potential to detect biomarkers of injury as well as characterize mechanisms of OA initiation and development[19, 20]. However, there are challenges to clinical implementation of synovial fluid biomarker analysis including technical difficulty and expense. Therefore, examination of serum metabolomic profiles may also yield biomarkers and phenotypic data for PTOA. Thus, the objective of this study is to examine metabolomic profiles of synovial fluid and serum in PTOA. To better understand the initiation of PTOA, we characterize changes in metabolomic profiles at early time points after injury. Toward developing serum biomarkers of PTOA, we examine correlations between metabolites co-detected in both the serum and synovial fluid. These results provide greater understanding of both the local and systemic response to joint injury and support further development of metabolomic biomarkers for OA as well as potential therapeutic targets.

## Materials and methods

### Preclinical Mouse Model of PTOA

Adult female C57BL/6 mice (n=12) were purchased from Charles River Labs and housed in the Animal Resource Center at Montana State University. After a two-week quarantine, mice were subjected to an established injury model[8, 9] at 12 weeks of age. Their care followed NIH guidelines and the protocol was approved by the Montana State University Institutional Animal Care and Use Committee.

Mice were randomly assigned to either control or injury groups and marked on their tails for identification. Of the 12 mice, 8 were injured while 4 served as uninjured controls. For injured mice, the injured stifle (*i.e.,* knee) joint (right or left) was randomized. The experimental sequence was the same for each mouse. Before injury, each mouse was placed in a chamber with 4% isoflurane and 2L/min O_2_ until anesthetized. Mice were then transferred to a nose cone with 3% isoflurane and 1L/min O_2_. All mice (injured and control) were then placed into the compression apparatus in a prone position.

The non-invasive injury apparatus consisted of a mechanical testing machine (Test Resources 510LE2, Shakopee, MN) outfitted with custom fixtures designed to position the hindlimb for stifle joint compression. The randomly selected stifle was placed in the cup with the ankle positioned directly above the knee in an ankle holder that provided ~30° of ankle dorsiflexion. A small pre-load (~1 N) stabilized the limb. For the mice in the injured group, a single dynamic axial compression at 130 mm/s with a target of 12N was applied. This caused anterior translation of the tibia in relation to the femur. After each compression, injury was confirmed by comparing anterior-posterior laxity to the contralateral joint. At high displacement rates as used here, this procedure results in mid-substance rupture of the ACL and leads to joint damage[8, 9, 21–23].

After injury, while still under anesthesia, all mice were administered 0.04 mL buprenorphine SR subcutaneously and monitored in a single-animal recovery cage until conscious and ambulatory. After recovery, demonstrated by walking in the cage, each mouse was placed in a fresh cage with its previous cage-mates. Control mice went through the exact same procedure as the injured mice including anesthesia, pre-load, and buprenorphine injection without the applied joint overload.

### Serum and Synovial Fluid Sample Collection

To examine early time point dynamics of PTOA, n=4 samples were harvested at 1- and 8-days post-injury along with n=2 controls at each time point which were pooled for analysis of n=4 controls. Blood was obtained by facial vein stick and collected in a standard red top serum tube microtainer (BD Microtainer Red Tubes No Additive, part number 365963). Approximately 200 μL of blood was collected for each mouse, and tubes were then inverted gently six times to begin activating the clotting factors. Each mouse was then immediately euthanized via cervical dislocation to avoid altering the blood pH through CO_2_ induced acidification. After allowing 30-60 minutes for activation of clotting factors, serum samples were centrifuged at 1,100 x g for 20 minutes to remove cells and clotting factors. This procedure was repeated for each mouse at their predetermined time point.

Synovial fluid was then extracted using established methodology[24]. Skin was removed from the injured leg and the tibial plateau was palpated with a scalpel to identify the joint line. The synovial membrane was accessed with an anterior incision through the patellar tendon that was then retracted proximally. A second incision was made through the joint capsule at which point, a 2 mm circular piece of Melgisorb (Tendra, part number 250600; Goteborg, Sweden) was dabbed onto the articular surfaces to adsorb synovial fluid. Once saturated, the Melgisorb was placed into a 1.5 mL microcentrifuge tube containing 35 μL of alginate lyase (Flavobacterium, Sigma-Aldrich A1603-100MG) in High Performance Liquid Chromatography (HPLC) water (1 U/mL concentration). Melgisorb was then digested in a water bath at 34°C for 30 minutes. To chelate calcium ions, 15 μL of 1.0M sodium citrate in HPLC water were added. After ~45 minutes of room temperature digestion, serum samples were centrifuged at 1100 x g for 15 minutes at 4°C. The supernatant was collected and transferred into a 1.5 mL microcentrifuge tube. All reagents and plasticware were HPLC-grade.

### Metabolite Extraction

To extract metabolites, serum and synovial fluid samples followed the same extraction protocol. Both were centrifuged at 500 x g for 5 minutes at 4°C to remove cells and debris. The supernatant was collected and mixed with 80% methanol (80:20 MeOH:H_2_O, vol/vol) at a ratio of 3:1 (weight:weight) methanol mixture:supernatant. This mixture was kept at −20°C for 30 minutes to precipitate macromolecules. Samples were then vortexed for 3 minutes and centrifuged at 16,100 x g for 5 minutes at 4°C. The supernatant was again removed and transferred to a new 1.5 mL microcentrifuge tube before being dried in a vacuum concentrator. Lipids and waxes were eliminated by re-extracting with 250 μL of an aqueous acetonitrile solution (1:1 acetonitrile:water, vol/vol) at 0°C for 30 minutes. They were again centrifuged at 16,100 x g for 5 minutes at 4°C and the supernatant was collected and dried in a vacuum concentrator. Samples were resuspended with 100 μL of aqueous acetonitrile solution (50:50 acetonitrile:water, vol/vol). They were vortexed for 3 minutes and 50 μL of each sample was transferred to a mass spectrometry tube and stored at −20°C until analysis.

### Metabolomics and Data Analysis

Metabolomic profiling was performed as previously described[19, 20] using LC-MS (liquid chromatography coupled to mass spectrometry). Chromatography used an Agilent 1290 UPLC system (Agilent, Santa Clara, CA, USA) and mass spectrometry was on an Agilent 6538 Q-TOF mass spectrometer (Agilent Santa Clara, CA, USA) in positive mode. The chromatography used a Cogent Diamond Hydride HILIC 150 x 2.1 mm column (MicroSolv, Eatontown, NJ, USA) in normal phase with established elution methods[19]. Spectra were analyzed as previously described[19]. This study used methods previously demonstrated to be effective in identifying possible biomarkers of OA. Raw mass spectrometry data files initially processed using XCMS[25] for normalization, alignment, and peak picking.

Data were analyzed in Metaboanalyst[26] using four groups: day 1 serum, day 1 synovial fluid, day 8 serum, and day 8 synovial fluid. Processing included normalization, log-transformation, and scaling to remove noise associated with mass spectrometry. Analyses assessed potential metabolic differences between the day 1, day 8, and control profiles for the serum and synovial fluid independently.

We used multivariate statistical methods to analyze metabolomic profiles to determine similarities and differences between serum and synovial fluid, as well as patterns of co-regulated metabolites. The overall variation in the dataset was assessed using unsupervised statistical methods including hierarchical cluster analysis (HCA) based on Euclidean distances and principal component analysis (PCA). To visualize the PCA, data were plotted as scatterplot projections on the principal axes. To assess differentially regulated metabolites, we used two-tailed Student’s t-tests with FDR corrections and volcano plot analysis. For metabolites co-detected between the serum and synovial fluid, we assessed relationships using Pearson correlation coefficients.

Finally, we used our established metabolomic profiling methods[20, 27, 28] to identify metabolite clusters using the complete linkage function based on the median metabolite intensities of each group. From each cluster, pathways were assessed based on the clustered metabolites using the MS Peaks to Pathways function within Metaboanalyst. This analysis used a 10-ppm tolerance in positive mode searching mummichog database with a significance threshold of 0.05. The *mus musculus* pathway set was used from the BioCyc database[29]. All analysis completed using *a priori* significance levels of p_fdr_ = 0.05 or for pairwise comparisons, false discovery rate (FDR) corrected significance levels of α = 0.05.

## Results

### Overview

To examine both the local and systemic response to joint injury, we characterized metabolomic profiles of synovial fluid and serum for control and injured mice. We detected 2544 total metabolite features across both the synovial fluid and serum samples (Figure 1). Of these, 2544 were present in synovial fluid, 2543 in serum, and 2533 in both synovial fluid and serum. 11 metabolites were found only in synovial fluid while 10 were found only in serum. While most metabolites were co-detected in both the synovial fluid and serum, only 230 metabolites (~9.0%) were of similar intensity (p > 0.05) between synovial fluid and serum. Both principal components analysis and hierarchical clustering find major differences in metabolomic profiles between serum and synovial fluid (Figure 1B-C). Synovial fluid appears to capture differences in metabolomic profiles between injured mice at both days 1 and 8 whereas serum does not (Figure 1C).

**Figure 1:**
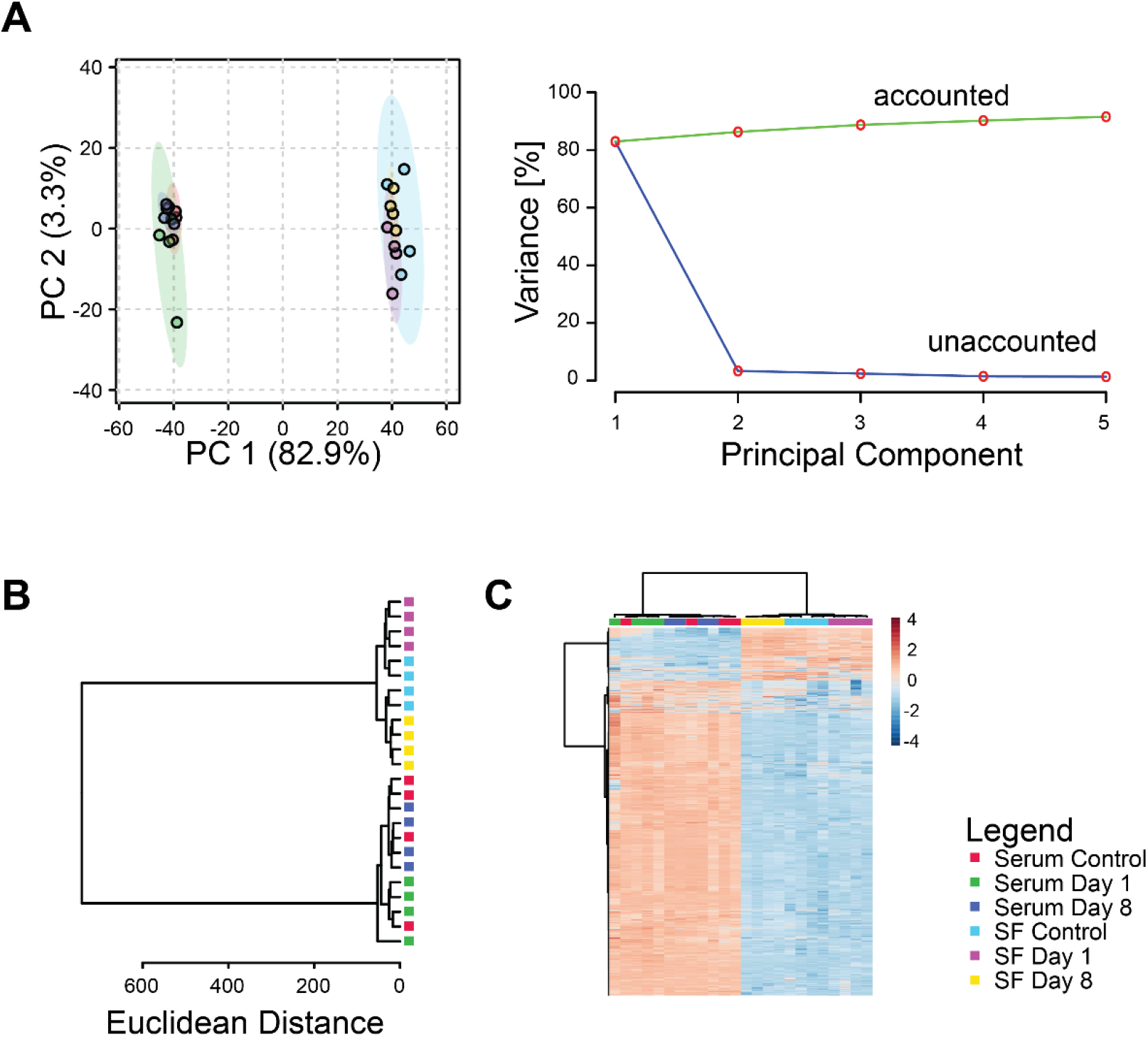
Differences in global metabolomic profiles between serum and synovial fluid. (A) Principal components analysis. Left projection of data onto the first 2 principal components shows distinct separation between metabolomic profiles of serum and synovial fluid. Right: scree plot shows that most of the variance in the overall dataset is associated with the first principal component. (B) Dendrogram of cluster analysis using the Euclidian distance metric and Ward linkage function shows complete separation between serum and synovial fluid samples. Furthermore, the synovial fluid metabolomic profiles were completely separated between experimental groups in contrast to the serum profiles that were not. (C) Heatmap of clustered metabolite intensities shows patterns of metabolite up- and down-regulation between experimental groups.

### Injury response within the Synovial Fluid

Metabolomic profiles of synovial fluid showed substantial differences between control and injured mice. Principal components analysis found distinct clustering between controls and injured samples and more than 50% of the overall variance was associated with the first three principal components (Figure 2A). Unsupervised cluster analysis found that both the day 1 and day 8 metabolomic profiles were distinct from controls (Figure 2B-C).

**Figure 2:**
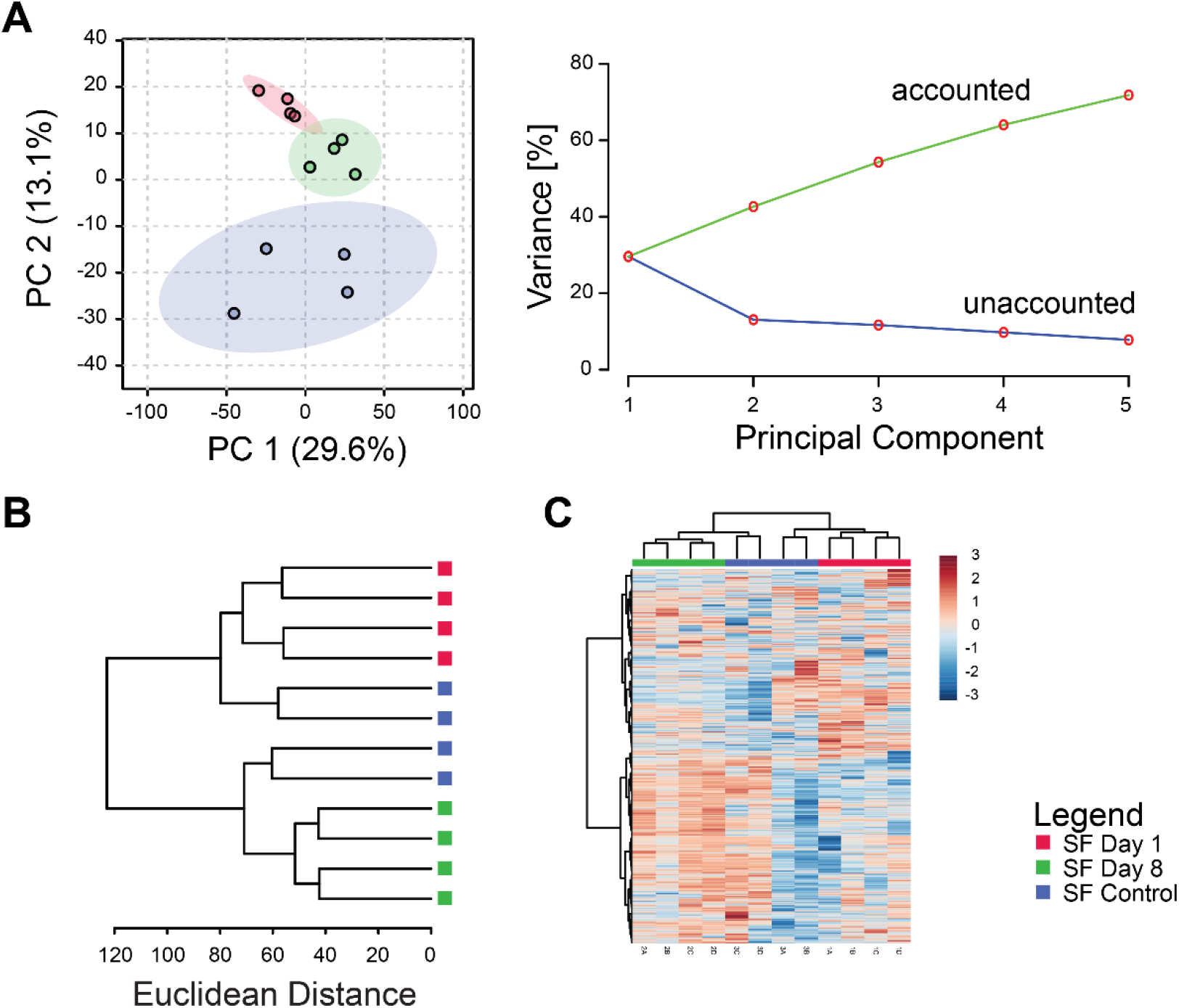
Synovial fluid metabolomic profiles capture changes induced by injury. (A) Principal components analysis. Left projection of data onto the first 2 principal components shows distinct separation between metabolomic profiles of control, 1-day post-injury, and 8-day post-injury synovial fluid. Right: scree plot showing that increasing the number of principal components results in increased modeled variance in the overall dataset. (B) Dendrogram of cluster analysis using the Euclidian distance metric and Ward linkage function shows complete separation between days 1 and 8 post injury. (C) Heatmap of clustered metabolite intensities shows patterns of metabolite up- and down-regulation between experimental groups.

### Injury response within the Serum

Serum metabolomic profiles were affected by joint injury, but less so than synovial fluid (Figures 2A and 3A). Principal components analysis found substantial overlap between controls and injured samples at day 8, but day 1 injured samples were more distinct (Figure 3A). The first three principal components were associated with 69% of the overall variance in the dataset. Unsupervised clustering failed to discriminate consistently between serum metabolomic profiles of controlled and injured mice (Figure 3B-C). Nonetheless, clusters B, D, F, and G (Figure 6) all demonstrated changes in the serum at different time points.

**Figure 3:**
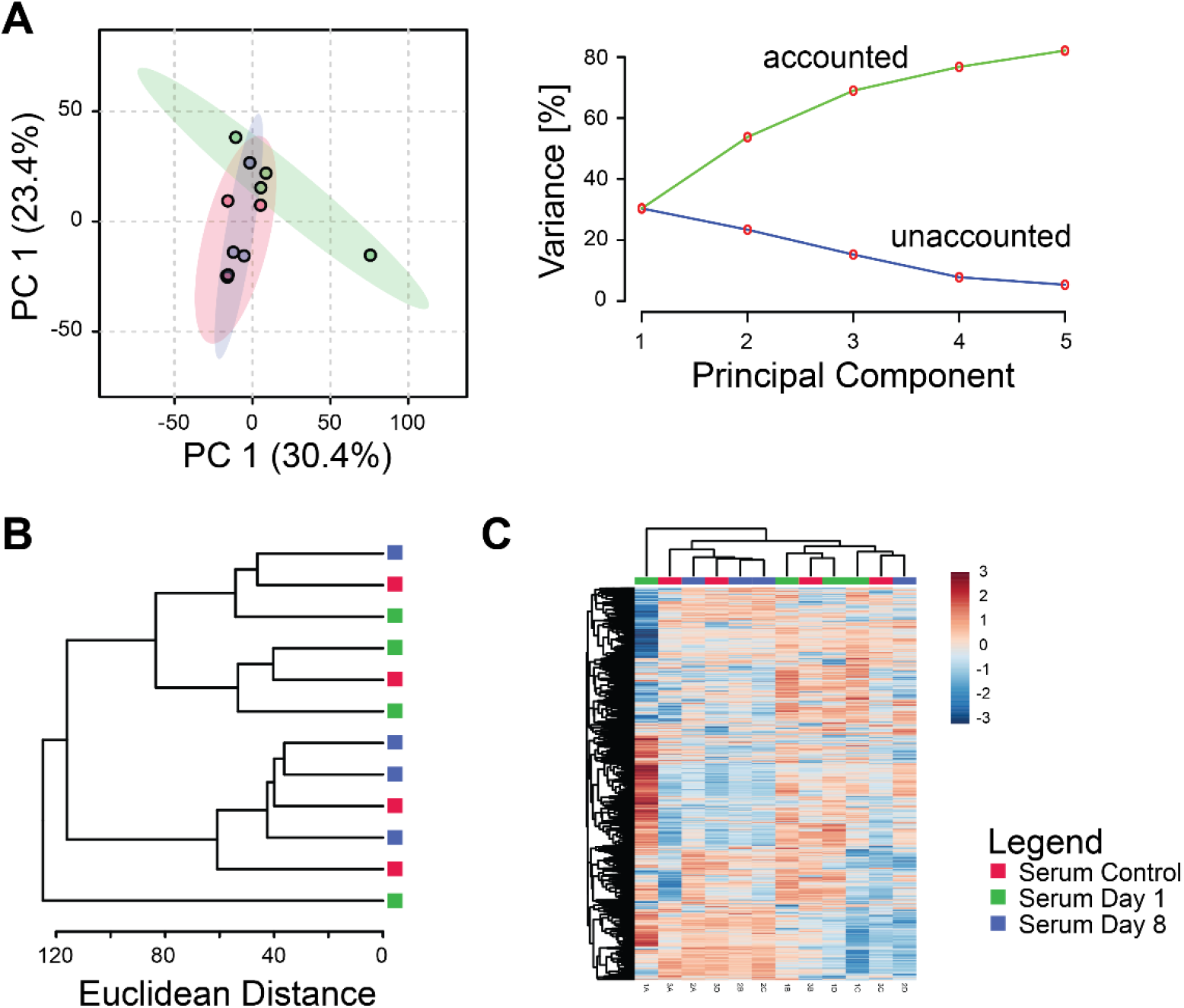
Injury response is partially captured using serum metabolomics. (A) Principal components analysis. Left: projection of data onto the first 2 principal components shows partial separation between serum metabolomic profiles of control, 1-day post-injury, and 8-day mice. Right: scree plot showing that increasing the number of principal components results in increased modeled variance in the overall dataset with a decreasing slope. (B) Dendrogram of cluster analysis using the Euclidian distance metric and Ward linkage function shows partial separation between experimental groups. (C) Heatmap of clustered metabolite intensities shows patterns of metabolite up- and down-regulation between experimental groups.

### Similarities and Differences between Metabolomic Profiles from Synovial Fluid and Serum

Metabolomic profiles capture responses to joint injury in both serum and the synovial fluid. However, there are substantial differences between metabolomic profiles of serum and synovial fluid at the same time point. At day 1 post-injury, PCA demonstrates clear separation between serum and synovial fluid (Figure 4A). Similar results are found using hierarchical clustering (Figure 4B-C), and volcano plot analysis shows many metabolites either up- or down-regulated in both groups, but the vast majority were upregulated in synovial fluid (Figure 4D). We found similar results when comparing synovial fluid and serum at day 8 post-injury (Figure 4E-H). These results indicate that distinct responses to injury occur between the synovial fluid and serum.

**Figure 4.**
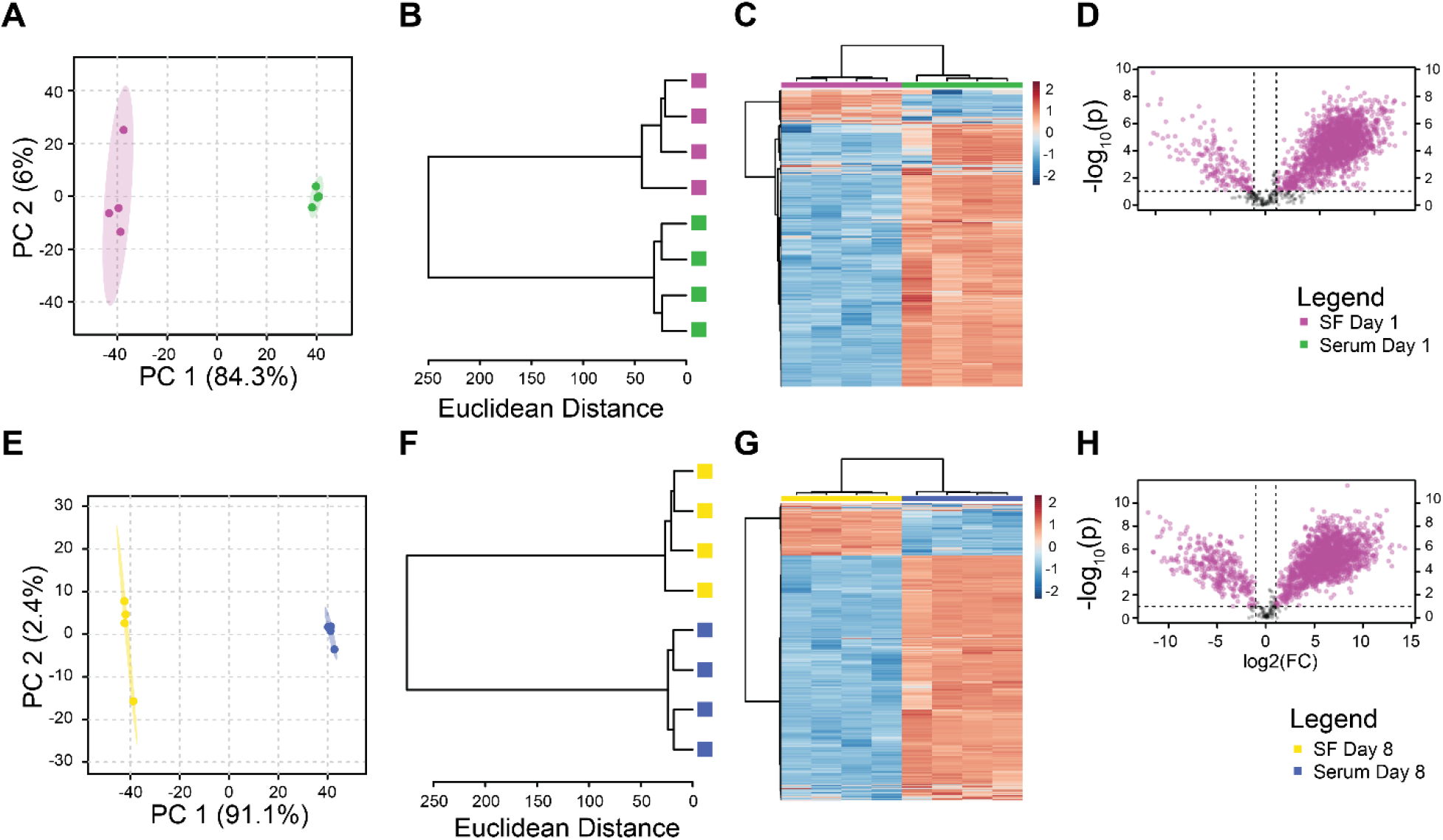
Paired analysis finds distinct metabolomic profiles between synovial fluid and serum at both day 1 and day 8 after injury. Day 1 panels A-D. Day 8 panels E-H. Principal components analysis (A, E). Unsupervised clustering (B, F). Clustered Heatmap (C, G). Volcano Plots (D, H).

While there are substantial differences in metabolite intensities, there were 2533 metabolites jointly detected in both serum and synovial fluid. To assess the potential for quantification of a synovial fluid metabolite level through measurement in the serum, we calculated correlation coefficients for the metabolite intensities between the serum and synovial fluid. There were 230 metabolite features with significant correlations between serum and synovial fluid (p < 0.05, Figure 5A). Of these correlated metabolites, 179 had positive correlations indicating that as the metabolite intensity increased in the serum it also increased in the synovial fluid. Of these positively correlated metabolite features, we identified 58 putative molecules including NADPH and steroid hormones (Figure 5B-C). 51 metabolites were negatively correlated indicating that increased intensity in the synovial fluid was associated with decreased intensity in the serum. Of these negatively correlated features, we identified 17 molecules including GAG monomers (Figure 5D). These significant correlations indicate that there is potential for using serum metabolite intensities to back-calculate synovial fluid metabolite intensities to provide insight into joint biology through assessment of serum metabolomic profiles.

**Figure 5.**
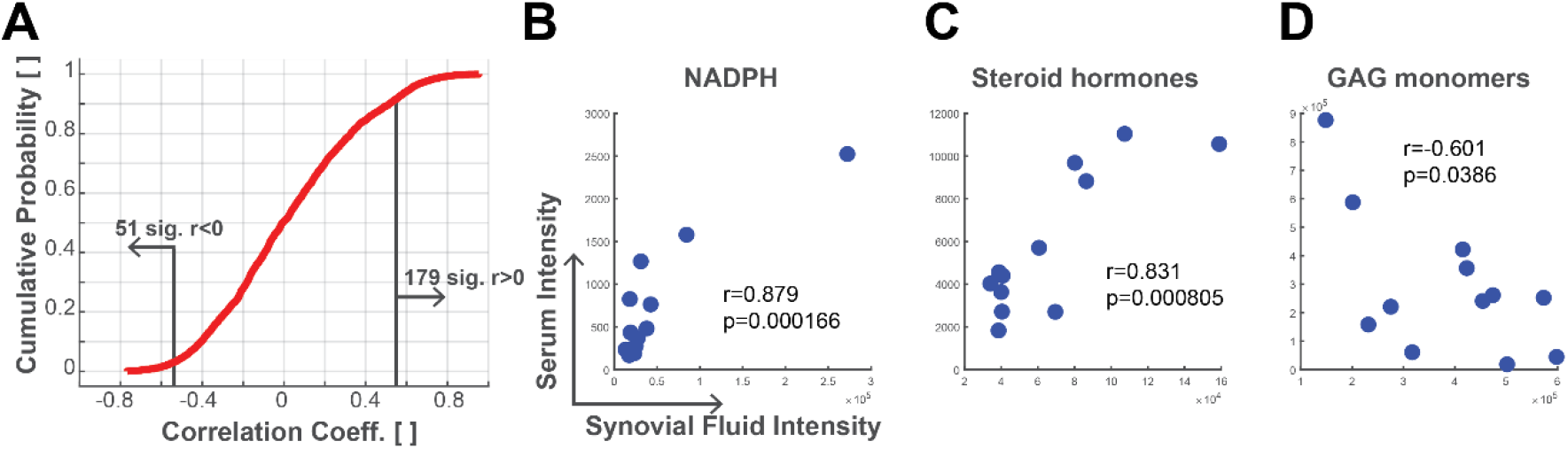
Correlations between synovial fluid and serum metabolites. Metabolite intensities were analyzed for correlations between serum and synovial fluid. (A) Cumulative probability plot of metabolite correlation coefficients. About 10% of co-detected metabolites have significant correlations (p<0.05). (B) Metabolite with putative identification of NADPH had a positive correlation between synovial fluid and serum. (C) Metabolite with putative database match for a steroid hormone including DHEA, testosterone, and androstenedione was positively correlated. (D) Metabolite with putative match representing the GAG monomers of N-acetyl-D-glucosamine and N-acetyl-D-galactosamine was negatively correlated.

### Pathways related to Joint Injury in the Synovial Fluid and the Serum

To assess pathways related to joint injury in each of the synovial fluid and serum compartments we identified clusters of co-regulated metabolites (Figure 6). In the synovial fluid (Figure 6A), cluster A was neutral in the control group, upregulated in day 1 post-injury, and downregulated in day 8 post-injury with no significant pathways. Cluster B was downregulated in the control group and upregulated in day 1 and day 8 post-injury, and also had no significant pathways. Cluster C was downregulated in the control group, upregulated in day 1 post-injury, and downregulated in day 8 post injury, mapping to asparagine degradation. Cluster D was upregulated in the control group and downregulated in day 1 and day 8 post injury, mapping to tryptophan degradation. Cluster E, which mapped to bupropion degradation, was neutral in the control group, downregulated in day 1 post-injury, and upregulated in day 8 post-injury. Cluster F mapped to tyrosine and phenylalanine degradation and was upregulated in the control group, downregulated in day 1 post-injury, and upregulated in day 8 post-injury. Cluster G was downregulated in the control group and day 1 post-injury but upregulated at day 8 post-injury. This cluster mapped to the tRNA charging pathway. Cluster H was also downregulated in the control group and day 1 post-injury and upregulated in day 8 post-injury but had no significant pathways.

**Figure 6.**
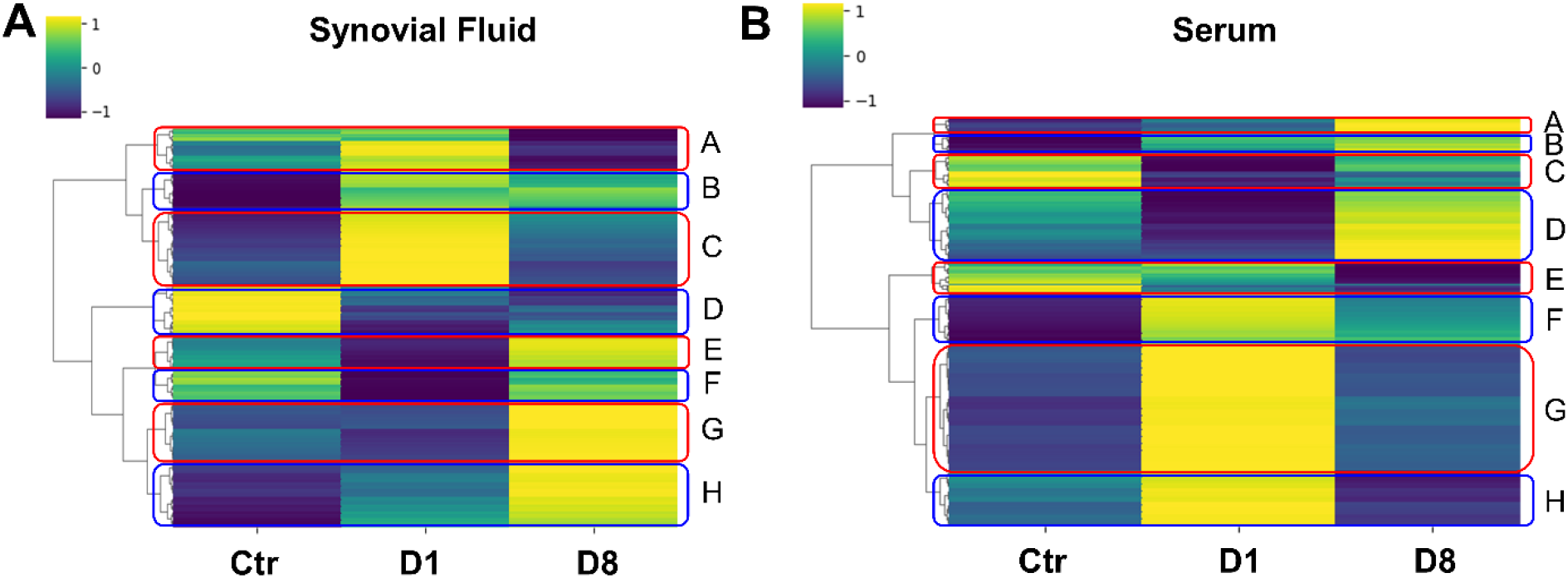
Clusters of coregulated metabolites within serum and synovial fluid indicate pathways relevant to joint injury. (A) Synovial fluid. (B) Serum. Median metabolite intensities for each group were subjected to clustering using the Euclidean distance metric. Clusters (A-H) of metabolites were then analyzed for pathways using Metaboanalyst.

In the serum (Figure 6B), cluster A was downregulated in the control group and day 1 post-injury and upregulated at day 8 post-injury but did not map to any significant pathways. Cluster B was downregulated in the control group, neutral in day 1 post-injury, and upregulated at day 8 post-injury, mapping to the tyrosine degradation pathway. Cluster C which had no significant pathways was upregulated in the control group, downregulated in day 1 post-injury, and neutral in day 8 post-injury. Cluster D was neutral in the control group, downregulated in day 1 post-injury, and upregulated at day 8 post-injury and mapped to the phospholipase pathway. Cluster E was upregulated in the control group, neutral in day 1 post-injury, and downregulated in day 8 post-injury with no significant pathways. Cluster F was downregulated in the control group, upregulated in day 1 post-injury, and neutral in day 8 post-injury mapping to arginine synthesis and proline degradation. Cluster G was downregulated in the control group, upregulated in day 1 post-injury, and downregulated at day 8 post-injury. This cluster mapped to the nicotine degradation pathway. Cluster H did not have any significant pathways and was downregulated in the control group, upregulated at day 1 post-injury, and downregulated in day 8 post-injury.

## Discussion

### Overview

This project provides insight to multiple different aspects of traumatic joint injuries and their effects on homeostasis. First, we gathered data regarding basic joint biology through synovial fluid immediately following traumatic joint injury while simultaneously collecting systemic data using serum. We were then able to compare the two in order to develop a complex understanding of local and systemic changes following joint injury. Lastly, we worked towards developing a biomarker panel for osteoarthritis.

### Injury Response within the Synovial Fluid

The synovial fluid samples demonstrated distinct changes in response to injury (Figure 2A). Five of the eight clusters (Figure 6A, clusters C, D, E, F, and G) related to pathways relevant to osteoarthritis initiation and progression.

At day 1 post-injury, cluster C was upregulated and mapped to the asparagine degradation pathway. Amino acids and their metabolites have been associated with end-stage osteoarthritis [30, 31] but it appears that they may play a role in acute injury as well[32]. However, this pathway became notably downregulated by day 8 post-injury indicating emphasis on an immediate response following injury.

At day 8 post-injury, clusters E, F, and G were upregulated. The pathway of interest in cluster E was bupropion degradation. This pathway most likely indicates metabolism of the buprenorphine given for pain relief during the experiment. Interestingly, this pathway was downregulated in day 1 post-injury before becoming upregulated at day 8 post-injury. However, the extended-release form which was given has a half-life of 43 to 60 days as described by the manufacturer which suggests why its metabolites were not found in the synovial fluid on day 1 post-injury. Both controls and experimental mice received the buprenorphine. In cluster F, the notable pathway mapped to tyrosine degradation. After a cell undergoes tyrosine hydroxylation it becomes catecholaminergic. One group showed that tyrosine hydroxylase positive cells were only present in the synovium of the patients with OA, not the controls[33]. It is thought that these catecholaminergic cells that release anti-inflammatory molecules may replace sympathetic nerve fibers which have grown into the cartilage over the progression of the disease and counteract the pro-inflammatory environment in the joint[34, 35]. Our findings show that while downregulated at day 1 post-injury, come day 8 post-injury, upregulation had occurred which may mark the initiation of catecholaminergic cells infiltrating the cartilage. The tRNA charging pathway was the notable pathway in cluster G. tRNA is a molecule that carries specific amino acids and matches them with mRNA to create proteins. Following an injury to the joint, protein production is increased to perform different functions within the cells. Our data suggests that immediately following the injury this process is downregulated but becomes upregulated at day 8 post-injury.

Only one cluster, cluster D, was upregulated in the control group. Cluster D mapped to the tryptophan degradation pathway which has been cited as a promising biomarker for osteoarthritis[36, 37]. Indoleamine 2,3-dioxygenase is responsible for degrading tryptophan into kynurenine and is induced by pro-inflammatory cytokines to counteract inflammation[32, 38]. The effect of indoleamine 2,3-dioxygenase and its role in inflammation is further being explored.

### Injury Response within the Serum

As a whole, the differences in serum between the control group, day 1 post-injury, and day 8 post injury were less distinct than in the synovial fluid (Figure 3A). However, four of the eight serum clusters (Figure 6B, clusters B, D, F, and G) showed changes in the serum across different time points. The pathways of interest for each cluster are discussed in the following paragraphs.

The clusters which were upregulated at day 1 post-injury included clusters F and G. Cluster F had metabolites mapping to synthesis of the amino acid arginine and degradation of the amino acid proline. Arginine has been indicated in inflammatory diseases given its role as an anti-inflammatory and has recently been shown to have decreased concentrations in patients with long-standing OA[30, 39, 40]. Arginine is also a precursor for proline which contributes to collagen and polyamine synthesis that results in fibrosis of tissue, further leading to OA[41]. Our data shows that these pathways were upregulated on day 1 post-injury which indicates that there is an initial increase in arginine synthesis and proline breakdown, both of which are protective in the short-term but damaging in the long-term. The metabolites of interest in cluster G mapped to the nicotine degradation pathway. Nicotine protects against joint inflammation and cartilage degradation in rodent models[42, 43]. The upregulation of nicotine degradation at day 1 post-injury in the serum indicates the protective role of such a pathway immediately following a traumatic injury. However, at day 8 post-injury, nicotine degradation became downregulated which may demonstrate that it provides only short-term anti-inflammatory properties.

At day 8 post-injury, clusters B and D had notably upregulated pathways. The metabolites found in cluster B mapped to the tyrosine degradation pathway which consists of catecholamines and melanin. Not only are catecholaminergic cells upregulated within the joint as mentioned previously, but also throughout the body, most notably in lymphoid organs. While doing an *in-vivo* mouse study using collagen-induced arthritis, it was observed that tyrosine hydroxylase positive cells were denser in draining lymph nodes, the thymus, and joints even prior to the onset of arthritis (days 5-21) indicating measurable changes prior to objective evidence of osteoarthritis[44]. Our data supports this notion in that this cluster was downregulated in the control group but became neutral in day 1 post-injury and upregulated in day 8 post-injury, indicating a gradual systemic catecholaminergic response to traumatic joint injury. Melanin, another byproduct of tyrosine, is the pigment that darkens our skin. Additionally, it has been found to be a potent anti-oxidative agent. Zhong et al. found that intra-articular injection of dopamine melanin nanoparticles sequestered numerous reactive oxygen species including hydroxyl radicals and superoxides. This was chondro-protective and led to decreased inflammatory cytokine release, decreased proteoglycan loss, and slower cartilage degradation when tested in rodent OA models undergoing ACLT surgery[45]. In cluster D, the metabolites of interest mapped to the phospholipase pathway which showed an initial downregulation prior to upregulation at day 8 post-injury. Phospholipase and its derivatives such as phosphatidylinositol-3-kinase (PI 3K) and phospholipase C (PLC) make up many of the cell signaling pathways including cell proliferation, intracellular trafficking, cell motility, cytoskeletal regulation, and ultimately cell death. Multiple studies have demonstrated that osteoblasts in joints with both PTOA and OA undergo modulated gene expression due to targeting of these pathways[46, 47].

### Similarities and Differences between Metabolomic Profiles from Synovial Fluid and Serum

Overall, there was great separation between synovial fluid and serum metabolites across all time points (Figures 1, 4). There was also distinct separation in synovial fluid between the control group and days 1 and 8 post-injury, while days 1 and 8 post-injury compared directly had slight overlap. The differences in serum were not as distinct, as all time points overlapped on PCA analysis. Paired analysis on day 1 post-injury showed that 84.3% variance between synovial fluid and serum was associated with the first principal component, and this increased to 91.1% on day 8 post-injury. Previous studies have also found that a minority of metabolites are correlated between the two fluids[48].

### Summary

In this study mice were subjected to non-invasive traumatic joint injury before sampling of synovial fluid and serum. Metabolomic profiles differed between these compartments with synovial fluid providing a more detailed description of injury-induced changes than serum. However, there were several metabolites correlated between these compartments indicating that a subset of information contained within the synovial fluid might be inferred through serum analysis for future development of biomarkers. Several key questions remain regarding joint injury including which cells and tissues are primarily characterized by synovial fluid analysis, potential differences between the male and female responses to injury, and the potential for translation of murine responses to humans.

## Acknowledgements

Funding provided by NSF (CMMI 1554708) and NIH (R01AR073964).

## Supplementary Information

Supplemental File 1 contains the metabolomics peak data used for processing in this study.

